# hb or not hb - when and why accounting for background mortality in toxicological survival models matters?

**DOI:** 10.1101/2023.01.25.525496

**Authors:** Julie Plantade, Virgile Baudrot, Sandrine Charles

## Abstract

Decisions in Environmental Risk Assessment (ERA) about impacts of chemical compounds on different species are based on critical effect indicators such as the 50% lethal concentration (*LC*_*50*_). Regulatory documents recommend concentration-response (or concentration-effect) model fitting on standard toxicity test data to get *LC*_*50*_ values. However, toxicokinetic-toxicodynamic (TKTD) models proved their efficiency to better exploit toxicity test data, at Tier-2 but also at Tier-1, delivering time-independent indicators. In particular, *LC*_*50*_ values can be obtained from the reduced General Unified Threshold model of Survival (GUTS-RED) with both variants, Stochastic Death and Individual Tolerance, that include parameter *h*_*b*_, the background mortality. Estimating *h*_*b*_ during the fitting process or not depends on studies and fitting habits, while it may strongly influence the other GUTS-RED parameters, and consequently the *LC*_*50*_ estimate. We hypothesized that estimating *h*_*b*_ from all data in all replicates over time should provide more precise *LC*_*50*_ estimates. We then explored how estimating *h*_*b*_ impacted: (i) GUTS-RED model parameters; (ii) goodness-of-fit criteria (fitting plot, posterior predictive check, parameter correlation); (iii) *LC*_*50*_ accuracy and precision. We finally show that estimating *h*_*b*_ does not impact the *LC*_*50*_ precision while providing more accurate and precise GUTS parameter estimates. Hence, estimating *h*_*b*_ would lead to a more protective ERA.

**Graphical abstract:** 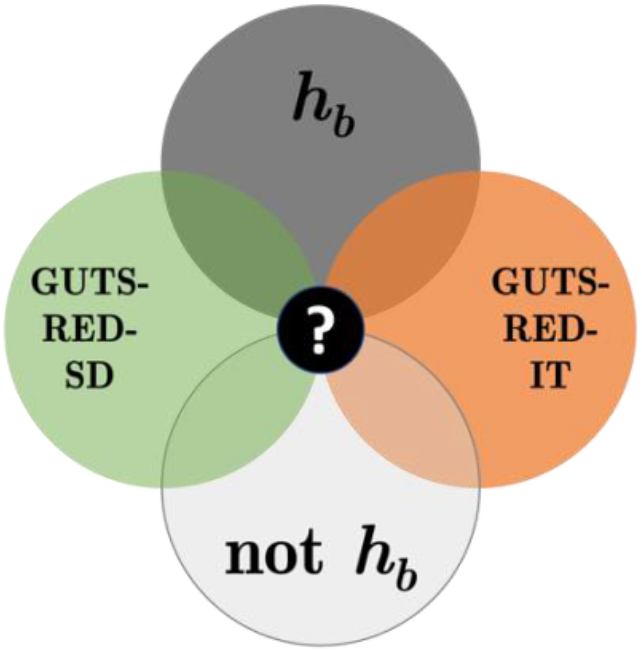

**Specifications table:** 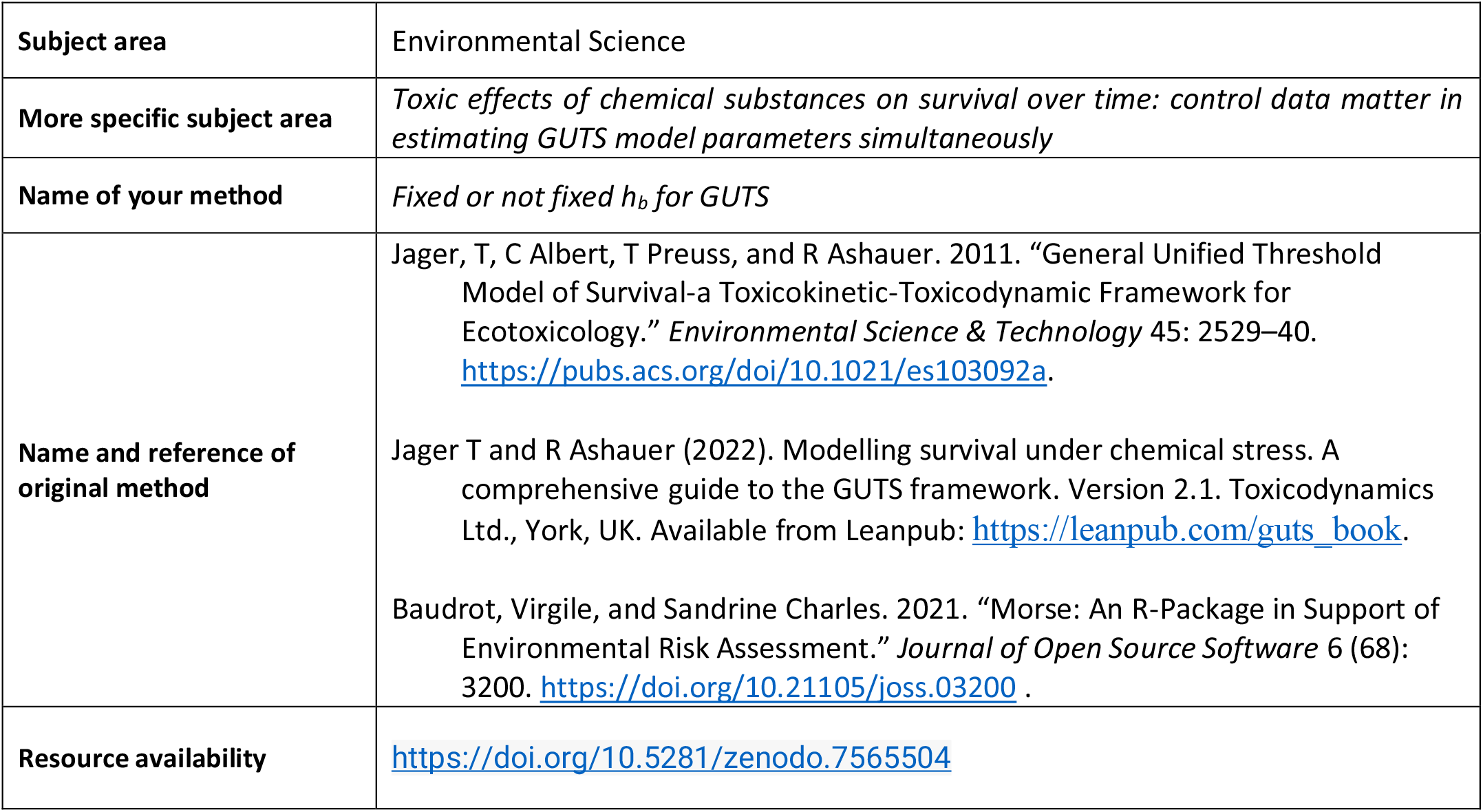

## Method details

### Motivation

Environmental Risk Assessment (ERA) aims at investigating the impact of chemical substances that can be potentially toxic, such as pharmaceuticals, plant protection products, metals, etc. This assessment is performed on different species, and it serves to advise regulatory bodies and professionals, for instance those of the food-processing industry. ERA is organized according to 4 tiers which extend from standard laboratory tests, on the one species / one compound principle, to ground tests at the ecosystem scale, including both the time and the space dimensions. Once Tier-1 is completed for the compound of interest on few classical species, several methods can be employed at Tier-2 to refine the effect potency evaluation [1]. Hence, in the perspective of better mimicking the environmental reality, additional toxicity tests can be performed under time-variable exposure profiles, either for the same species as those used at Tier-1, or for additional ones. Through use of toxicity test data at both Tier-1 and Tier-2, the environmental risk can be assessed by employing toxicokinetic-toxicodynamic (TKTD) models that allow to account for both the concentration- and the time-dependencies.

Recently, TKTD models have been recognized advantageous for the risk assessment, especially for aquatic organisms exposed to pesticides [2]. Among available TKTD models, the General Unified Threshold model of Survival (GUTS) was particularly highlighted being considered as ready-to-use with survival test data within a regulatory context. Indeed, the GUTS theoretical framework is well established [3, 4], and turnkey tools are available to implement it. On a general point of view, GUTS allows (i) to translate the exposure concentration over time into an internal dose that may cause damages, and (ii) to translate these damages in terms of visible effects on the individual survival over time. Once calibrated on standard survival test data, and validated on additional survival data (e.g., collected under time-variable exposure profiles or mixture toxicity tests), GUTS serves to predict lethal concentrations *LC(x,t)* at any target time *t* and for any chosen x% [5] thus providing refined toxicity indicators in support of decision makers.

In this paper, we focus on the context of regulatory ERA. As stated by EFSA [2], GUTS may be used in its reduced (RED) version, itself comprising two variants: the Stochastic Death (GUTS-RED-SD) variant and the Individual Tolerance (GUTS-RED-IT) variant. Both variants include four parameters, among them the toxicokinetic parameter *k*_*d*_, called the dominant rate constant; and the background mortality rate *h*_*b*_, corresponding to the natural mortality that may occur even if the organisms are not exposed to a chemical stressor. Because of a presumed independence from parameters directly related to the potential toxicity of the chemical of interest (i.e., *k*_*d*_ and the two others), the *h*_*b*_ parameter is often fixed from the control data, while the other three GUTS parameters are estimated by fitting the model to all of the collected data. This way of doing has pros and cons which this paper aims at investigating with regards to a simultaneous fitting of GUTS providing estimates of the four parameters.

Independently setting the *h*_*b*_ parameter before the fitting process means that only three of the four GUTS parameters will be estimated. Fixing the *h*_*b*_ parameter may be a biological choice (e.g., no individual died until the end of the experiment under control conditions), or a computing-based choice (e.g., either to facilitate the convergence of the fitting process, or to increase the estimation precision, when data are sparse or when the experiment was not ideally designed). However, setting the *h*_*b*_ parameter implies several implicit assumptions: (i) the *h*_*b*_ parameter is not correlated to the three other GUTS parameters, and (ii) there is no additional information on the background mortality within the toxicity data. Consequently, setting the *h*_*b*_ parameter amounts to neglect the fact that the experimental design may contain very low tested concentrations for which no effect can be observed, as under control conditions. Also, neglecting the potential correlation of *h*_*b*_ with the three other parameters may impact their respective estimates, and subsequently the *LC(x,t)* estimate.

So, the aim of this paper is to evaluate the impact of estimating or not the *h*_*b*_ parameter, on both the estimates of the three other GUTS parameters, and the subsequent estimation of the *LC*_*50*_ at final time. We performed our study with both variants of the reduced GUTS, namely GUTS-RED-SD and GUTS-RED-IT, on a collection of 38 data sets comprising different species-compound combinations. We first discussed the results from a modelling perspective, to statistically assess the impact of estimating or not estimating *h*_*b*_; then discussed in the perspective of regulatory ERA, to propose the best compromise between the rapid and daily completion of risk assessments and having sufficient scientific and statistical rigor so that the results on which decisions are based are reliable.

### Modelling

Both variants of the reduced GUTS, namely GUTS-RED-SD and GUTS-RED-IT, were systematically used, both under option 1 (*h*_*b*_ simultaneously estimated with the other three parameters) and option 2 (*h*_*b*_ estimated on control data only), as detailed below. The GUTS-RED-SD variant assumes that all individuals are identically sensitive to the chemical substance by sharing a common threshold internal dose, and that mortality is a stochastic process whose intensity is linearly increasing once this threshold is reached. In contrast, the GUTS-RED-IT variant is based on the critical body residue approach, which assumes that individuals differ in their thresholds following a given probability distribution and that they die as soon as the internal dose reaches the individual-specific threshold [5].

Option 1 (denoted O1) stands for the fitting of the two GUTS-RED variants in their complete form, both involving four parameters, among which *k*_*d*_ and *h*_*b*_ as described earlier. In addition, variant GUTS-RED-SD has parameters *z*_*w*_ (the internal threshold from which effects may appear) and *b*_*w*_ (the “killing rate”, that is a parameter related to the intensity of the effect) to describe the instantaneous mortality rate as a function of the internal dose via a linear stepwise function [4]:

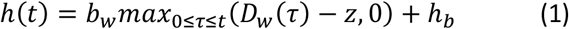

where *D*_*w*_(*τ*) stands for the internal dose at time *τ*. Variant GUTS-RED-IT also has two additional parameters, related to the fact that the individual threshold is usually assumed to follow a log-logistic probability distribution, of median *m*_*w*_ and spread β. This leads to the following cumulative distribution function:

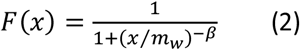

with *max*_0<*τ*<*t*_(*D*_*w*_(*τ*)) = *x*, the maximal internal dose reached during the exposure period.

Option 2 (denoted O2) stands for the fitting of the two GUTS-RED variants, the *h*_*b*_ parameter being fixed from control data only, according to the following calculation:

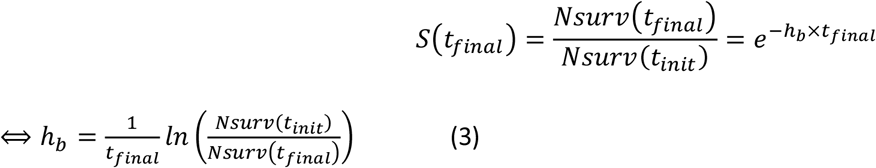

In equation (3), *Nsurv* stands for the observed number of survivors either at initial time, *Nsurv*(*t*_*init*_), or at final time, *Nsurv*(*t*_*final*_), of the experiment. In the end, four situations were investigated crossing the two GUTS-RED variants (SD and IT) with the two options (O1 and O2). Hereafter, each situation is always associated with the same color code: dark green for SD-O1; light green for SD-O2; dark orange for IT-O1; light orange for IT-O2. Both GUTS-RED variants were fitted with the R-package “morse” [6] under R version 4.2.1 (“Funny-Looking Kid”, 2022-06-23). In particular, function “survFit()” was used, with option “model_type” equal to “SD” (for variant GUTS-RED-SD) or “IT” (for variant GUTS-RED-IT) and option “hb_value” equal TRUE (for option 1) or FALSE (option 2). Under option 2, parameter *h*_*b*_ was always calculated before the fitting process. Finally, function “doseResponse_survFitCstExp” of package “morse” was adapted to get the posterior probability distribution of the *LC*_*5*0_ estimate for each of the four situations SD-O1, SD-O2, IT-O1 and IT-O2.

### Data sets

To ensure reliable results, the above-mentioned procedure was used with a total of 38 data sets (Table 1). Each data set was fitted under each of the four situations SD-O1, SD-O2, IT-O1 and IT-O2, leading to a total of 152 fitting outputs. For each data set, fitting outputs of SD and IT variants were stored in two separated PDF files, each including for both options 1 and 2: (i) fitting plots ; (ii) parameter estimates, expressed with the median, the 95% credible interval and the precision ; (iii) the fixed *h*_*b*_ value (option 2 only) ; (iv) the posterior predictive check (PPC) ; (v) the comparison of prior and posterior marginal probability distributions for each estimated parameter ; (vi) the correlation matrix between estimated parameters ; (vii) the deviance information criteria (DIC) and the calculation of the difference between option 1 and option 2. Note that a negative value of the “Diff_DIC” indicates that the model with parameter *h*_*b*_ simultaneously estimated with the three other parameters will be preferred. Finally, *LC*_*50*_ estimates are given for both options 1 and 2, expressed with the median and the 95% credible interval. All these output files were analyzed regarding the different goodness-of-fit criteria: (i) the visual fit, (ii) the PPC, (iii) the prior-posterior comparison, and (iv) the correlations between parameters.

**Table 1:** Details on the 38 data sets that were analyzed in this study, namely information about the species and the chemical compound used for toxicity tests for the 38 data sets.

| Code name | Species | Chemical | Comments | Reference |
| --- | --- | --- | --- | --- |
| TOX03-SPE19 | *Daphnia magna* | Copper |  | [7] |
| TOX04-SPE19 | *Daphnia magna* | Zinc |  | [7] |
| TOX10-SPE19 | *Daphnia magna* | Cadmium |  | [7] |
| TOX10-SPE15 | *Lymnaea stagnalis* | Cadmium |  | [8, 9] |
| TOX05-SPE19 | *Daphnia magna* | Dichromate potassium |  | [10] |
| TOX06-SPE19 | *Daphnia magna* | Oxolinic acid |  | [10] |
| TOX07-SPE19 | *Daphnia magna* | Streptomycine |  | [10] |
| TOX08-SPE19 | *Daphnia magna* | Sulfadiazine |  | [10] |
| TOX12-SPE16 | *Pimephales promelas* | NA |  | [11] |
| TOX01-SPE02_b | *Asellus aquaticus* | Chlorpyrifos |  | [12] |
| TOX01-SPE03_b | *Chaoborus crystallinus* | Chlorpyrifos |  | [12] |
| TOX01-SPE04_b | *Cloeon dipterum* | Chlorpyrifos |  | [12] |
| TOX01-SPE17 | *Epeorus assimilis* | Chlorpyrifos |  | [12] |
| TOX01-SPE18 | *Rhithrogena semicolorata* | Chlorpyrifos |  | [12] |
| TOX01-SPE19-11 | *Daphnia magna* | Chlorpyrifos | T = 11°C | [12] |
| TOX01-SPE19-14 | *Daphnia magna* | Chlorpyrifos | T = 14°C | [12] |
| TOX01-SPE19-17 | *Daphnia magna* | Chlorpyrifos | T = 17°C | [12] |
| TOX01-SPE19-20-1 | *Daphnia magna* | Chlorpyrifos | T = 20°C, n°1 | [12] |
| TOX01-SPE19-20-2 | *Daphnia magna* | Chlorpyrifos | T = 20°C, n°2 | [12] |
| TOX09-SPE19 | *Daphnia magna* | Ivermectine |  | [13] |
| TOX02-SPE19 | *Daphnia magna* | Chlordan |  | [14] |
| TOX11-SPE05 | *Gammarus pulex* | Propiconazole |  | [15] |
| TOX01-SPE01 | *Anax imperator* | Chlorpyrifos |  | [16] |
| TOX01-SPE02_a | *Asellus aquaticus* | Chlorpyrifos |  | [16] |
| TOX01-SPE03_a | *Chaoborus obscuripes* | Chlorpyrifos |  | [16] |
| TOX01-SPE04_a | *Cloeon dipterum* | Chlorpyrifos |  | [16] |
| TOX01-SPE05 | *Gammarus pulex* | Chlorpyrifos |  | [16] |
| TOX01-SPE06 | *Neocaridinia denticulata* | Chlorpyrifos |  | [16] |
| TOX01-SPE07 | *Notonecta maculata* | Chlorpyrifos |  | [16] |
| TOX01-SPE08 | *Parapoynx stratiotata* | Chlorpyrifos |  | [16] |
| TOX01-SPE09 | *Plea minutissima* | Chlorpyrifos |  | [16] |
| TOX01-SPE10 | *Procambarus spec. (adults)* | Chlorpyrifos |  | [16] |
| TOX01-SPE11 | *Procambarus spec. (neonates)* | Chlorpyrifos |  | [16] |
| TOX01-SPE12 | *Ranatra linearis* | Chlorpyrifos |  | [16] |
| TOX01-SPE13 | *Sialis lutaria* | Chlorpyrifos |  | [16] |
| TOX01-SPE19 | *Daphnia magna* | Chlorpyrifos |  | [16] |
| A1 | Simulated with GUTS-RED-SD, true $h_b$ value = 0.01 | | | [4] |
| A2 | Simulated with GUTS-RED-IT, true $h_b$ value = 0.02 | | | [4] |

### Outcomes

In general, our results show that option 1 (that is estimating the *h*_*b*_ parameter) provides more precise and accurate estimates of the three other parameters for both SD and IT GUTS-RED variants. Surprisingly, our results show that simultaneously estimating the four parameters of GUTS-RED variants does not provide better *LC*_*50*_ estimates. Consequently, in terms of ERA, if only the *LC*_*50*_ is of interest, fitting GUTS-RED variants by fixing *h*_*b*_ (option 2) could be sufficient.

### Influence of estimating *h*_*b*_ on parameter estimates

As illustrated in Figure 1, for both data sets A1 and A2 (Table 1), the estimated *h*_*b*_ values under option 1 (*h*_*b,1*_) are closer to the true *h*_*b*_ value than the calculated *h*_*b*_ value under option 2 (*h*_*b,2*_). This results in favor of option 1 is also observed with the other data sets we analyzed (see supplementary information). For all data sets except one, whatever the GUTS-RED variant, values of *h*_*b,1*_ and *h*_*b,2*_ were different; the exception was obtained with data set TOX01-SPE12 (Table 1, Figure S1), a particular data set with only four time points and a very wide range of tested concentrations that could explain some fitting difficulties whether the *h*_*b*_ parameter is estimated or not. As also illustrated in Figure S1, option 2 tends to underestimate the background mortality, with 56/72 *h*_*b,1*_ estimates greater than *h*_*b,2*_ values. In addition, for all datasets, *h*_*b,1*_ estimates are quite precise, both with SD and IT. Even though it may seem obvious, estimating parameter *h*_*b*_ with the others allow the quantification of its uncertainty in the same way.

**Figure 1:**
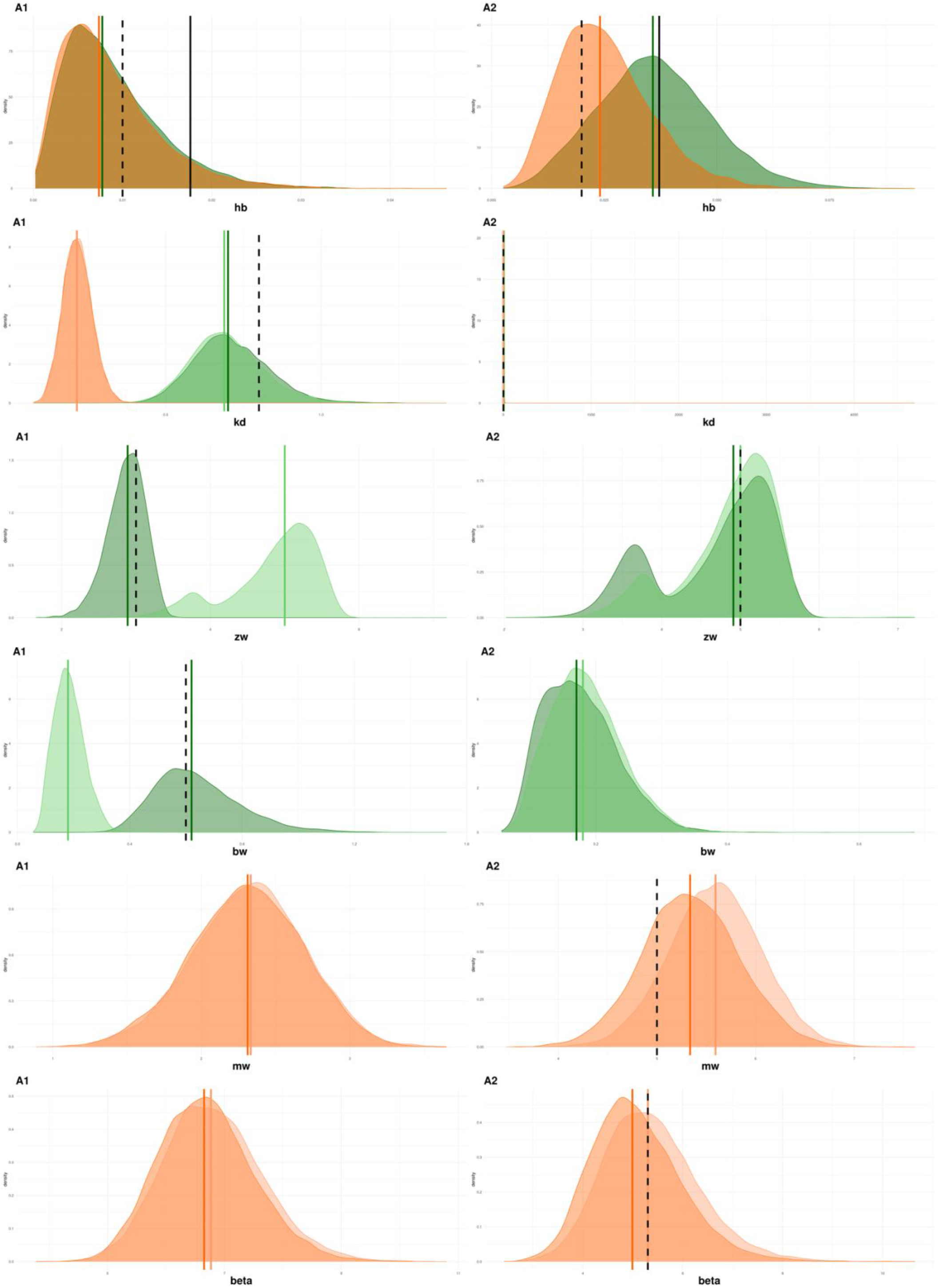
Marginal posterior probability distributions of both GUTS-RED variants, SD (in green), IT (in orange), for both options 1 (dark colors) and 2 (light colors), and for data sets A1 and A2. Colored solid vertical lines stand for the median of the posterior probability distribution; the black dashed vertical line stands for the true parameter values as used to simulate data points of data sets A1 and A2; and the black solid vertical line stands fo the calculated fixed h_b_ value.

Regarding the other parameters, Figure 1 shows that, under option 1, *z*_*w*_ and *b*_*w*_ (data set A1), as well as *m*_*w*_ (data set A2) are closer to the true value than under option 2. Firstly, option 2 overestimates *z*_*w*_ with GUTS-RED-SD (data set A1) and *m*_*w*_ in GUTS-RED-IT (data set A2). This result for *z*_*w*_ is confirmed with the remaining data sets, among which 14/36 data sets show a difference in *z*_*w*_ estimation between option 1 (*z*_*w,1*_) and 2 (*z*_*w,2*_, Figure S3). Twelve out of these 14 data sets lead to a *z*_*w,1*_ value greater than the *z*_*w,2*_ value. The two exceptions concern data sets TOX01-SPE17 and TOX01-SPE18 with a long exposure period of 96 days during which 13 time points were distributed in triplets of closely spaced time points. Also, in these two data sets, the range of the tested concentrations is very large with quite widely spaced concentrations which could prevent a precise estimate of *z*_*w*_ whatever the option. Similar results were obtained for *m*_*w*_ with GUTS-RED-IT. Indeed, seven data sets show different *m*_*w*_ estimates between option 1 (*m*_*w,1*_) and 2 (*m*_*w,2*_, Figure S5). Six out of these seven data sets show a *m*_*w,1*_ greater than the *m*_*w,2*_ value, the only exception being obtained with data set TOX01-SPE17 for which difficulties have already been identified (see above) for the estimation of *z*_*w*_. Because parameters *z*_*w*_ and *m*_*w*_ correspond to a No-Effect Concentration (NEC) within the model, this means that option 2 tends to overestimate the internal dose above which the chemical compound will have an effect on individuals.

Secondly, whatever the data sets, our results reveal that option 2 underestimates *b*_*w*_ in GUTS-RED-SD (see Figure 1, for an illustration with data sets A1 and A2). Ten data sets led to *b*_*w*_ estimates different between options 1 (*b*_*w,1*_) and 2 (*b*_*w,2*_, Figure S4). Out of these 10 data sets, eight data sets led to *b*_*w,2*_ less than *b*_*w,1*_. The two other data sets are TOX01-SPE03_a, and TOX05-SPE19 for which the dose-response curve at final time is very steep, making difficult the estimation of *b*_*w*_ whatever the option. So, reminding that parameter *b*_*w*_ stands for the intensity of the chemical effect on individuals, it appears that option 2 tends to underestimate this effect.

While the β parameter does not inspire any specific comment, the *k*_*d*_ parameter that corresponds to the toxicokinetic part of the model, also appears close to the true value but only with GUTS-RED-SD, indeed GUTS-RED-IT clearly underestimates *k*_*d*_ (Figure 1, data set A1). Similar results were obtained with the other data sets. A total of 18/36 data sets fitted with both GUTS-RED-SD and GUTS-RED-IT gave different *k*_*d*_ estimates whatever the option (Figure S2). For these 18 data sets, the *k*_*d*_ estimate with GUTS-RED-IT was systematically lower than the one obtained with GUTS-RED-SD. This means that for half of the data sets we analyzed, the speed of the kinetic is underestimated, either by GUTS-RED-SD or GUTS-RED-IT. This corroborates the current well-known inability to anticipate the use of one or the other GUTS-RED variants for a particular species-compounds combination [2].

Our results on parameter estimates lead us to the conclusion that estimating the background mortality (option 1) provides *h*_*b*_ values closer to the reality. Because option 2 tends to overestimate *z*_*w*_ and *m*_*w*_, and to underestimate *b*_*w*_, using option 1 will provide more protective predictions in terms of ERA.

### Influence of estimating *h*_*b*_ on goodness-of-fit criteria

Looking at fitting plots of posterior predictive checks for all data sets shows no difference whether parameter *h*_*b*_ is estimated or not. Looking at parameter correlations seems to be the most convenient way to highlight the impact on estimating *h*_*b*_ or not on the estimation of the three other parameters. Indeed, fixing *h*_*b*_ before the inference process leading the other three estimates (option 2) means completely ignoring any correlation of *h*_*b*_ with the other three parameters. However, as illustrated in Figure 2, not estimating *h*_*b*_ hardly changes correlations between the other three parameters. In addition, we can see that when estimated, *h*_*b*_ is only weakly correlated with the other three parameters. This result holds true for all other datasets we analyzed. Finally, whatever the GUTS-RED variants we used, whatever the datasets we analyzed, whatever the goodness-of-fit criteria we looked at, there was virtually no difference between the options 1 and 2. Nevertheless, because the parameters are better estimated under option 1 (see above-mentioned arguments and Figure 1), we discuss only based on option 1 hereafter.

**Figure 2:**
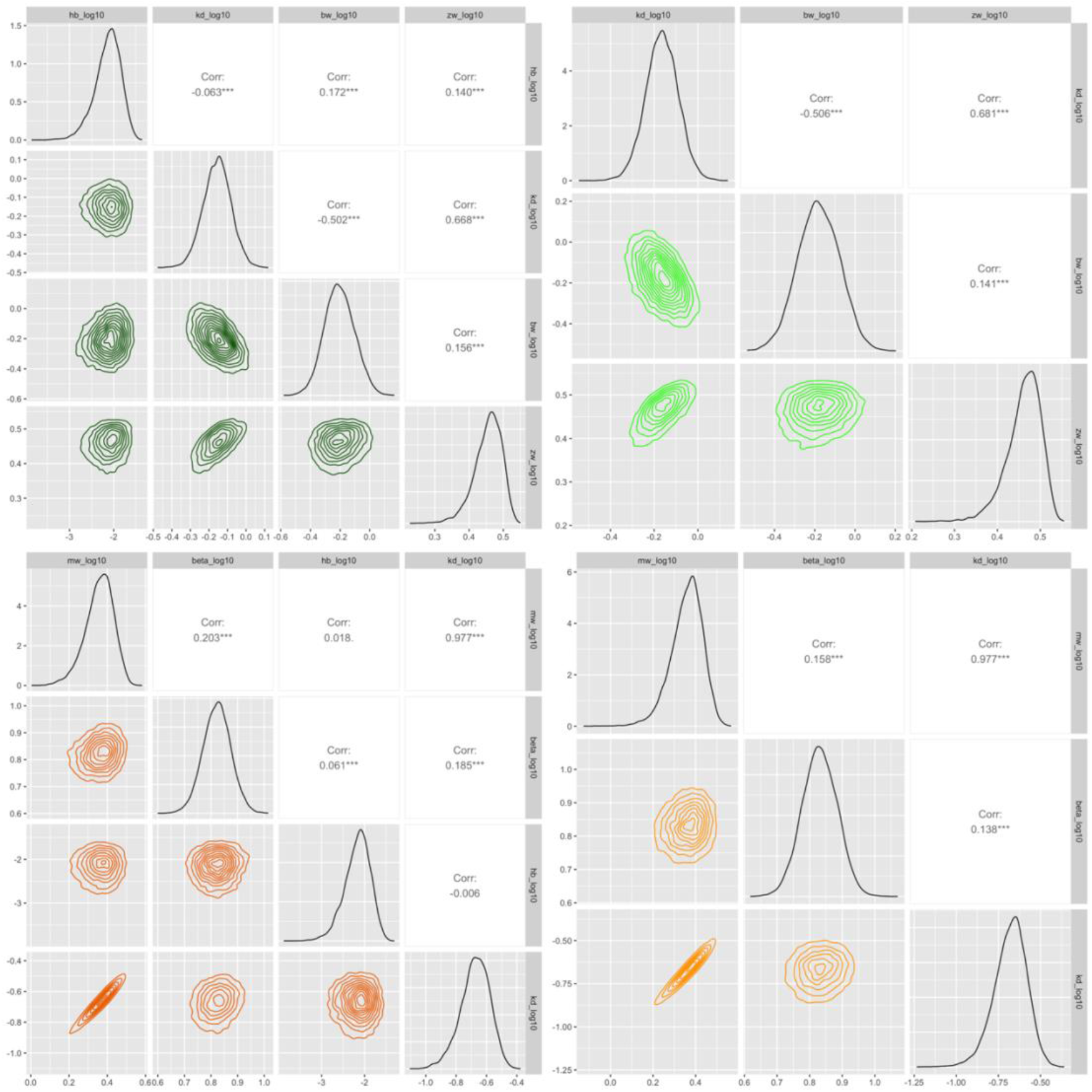
parameter correlations for data set A1: upper left with SD-O1; upper right with SD-O2; lower left with IT-O1; lower right with IT-O2.

Figure 3 confirms that *h*_*b*_ is poorly correlated to the other three parameters for all data sets. Correlation coefficients rarely exceed 0.25, and when exceeding 0.25, it happens more often with GUTS-RED-IT than with GUTS-RED-SD. With the exception of two or three data sets, parameters of the GUTS-RED-IT variant are always positively correlated, while correlation coefficients are distributed around 0 with GUTS-RED-SD. The *h*_*b*_ parameter is most often highly correlated with the *β* parameter, and the correlations of *h*_*b*_ with both *k*_*d*_ and *m*_*w*_ is very similar most of the time; this is due to a strong correlation between *k*_*d*_ and *m*_*w*_ that inherently exists in GUTS-RED-IT [18]; see Figure S8 to visualize this for all the data sets we analyzed in this study. With variant GUTS-RED-SD, the correlation coefficients are more often negative between the parameters *h*_*b*_ and *k*_*d*_ than with the two other parameters; also, when positive, this correlation is very small. Biologically speaking, this means that the background mortality rate is more correlated to the toxicokinetic than to the toxicodynamic of the compound. Such a result is particularly interesting because it highlights a possible trade-off between the survival of organisms and their bioaccumulation of chemical substances.

**Figure 3:**
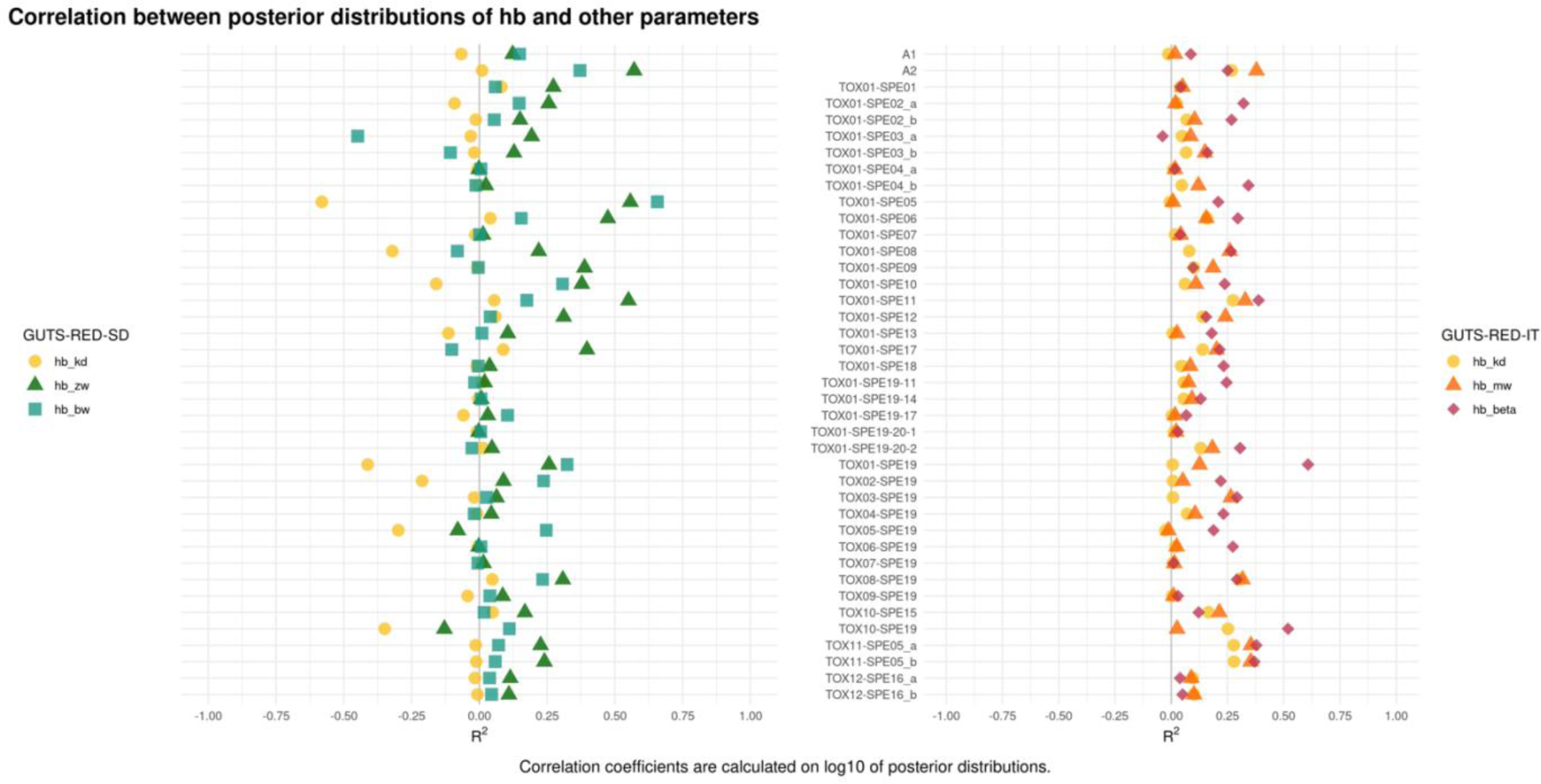
parameter correlations between parameter h_b_ and the three other ones for all data sets and both GUTS-RED-SD (left panel) and GUTS-RED-IT (right panel).

### Influence of estimating *h*_*b*_ on the *LC*_*50*_ estimate

As illustrated on Figure 4, whatever the data set, with one or two exceptions, there is almost no difference between the *LC*_*50*_ with both GUTS-RED-SD and GUTS-RED-IT whether parameter *h*_*b*_ is estimated or not. However, *LC*_*50*_ estimates may sometimes strongly differ whether GUTS-RED-SD or GUTS-RED-IT is used (see for example Figure 4, on the right). Two *LC*_*50*_ estimates were considered significantly different when their 95% credible intervals did not or very few overlap. Hence, in total, only 6 out of the 38 data sets exhibited significantly different *LC*_*50*_ values. This result is consistent with other works also illustrating the impossible choice between both GUTS-RED variants [2].

**Figure 4.**
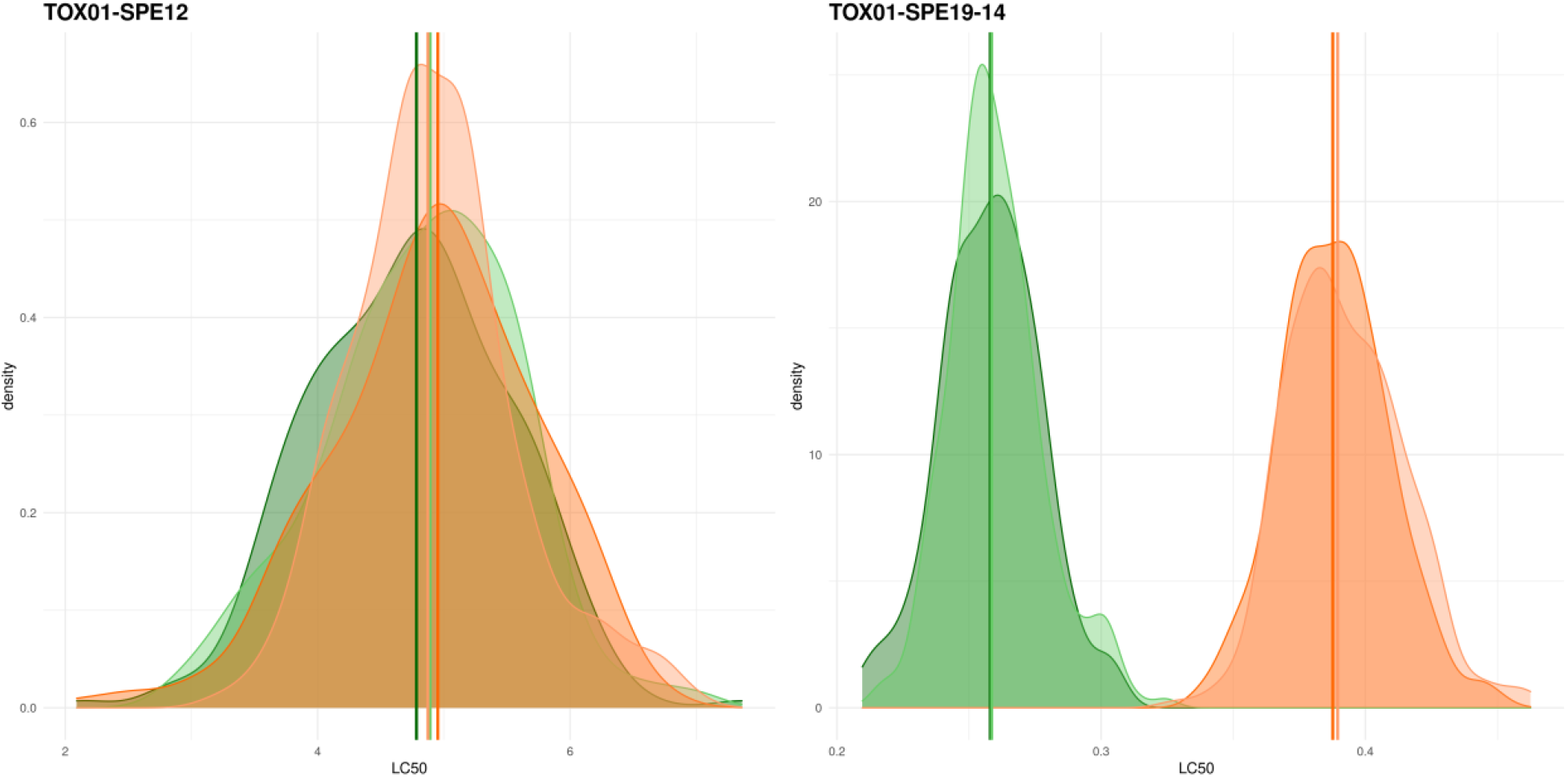
Distributions of LC_50_ at final time with SD O1 (dark green), SD O2 (light green), IT O1 (dark orange) and IT O2 (light orange): on the left for data set TOX01-SPE12; on the right for data set TOX01-SPE19-14. See table 1 and 2 for details on data sets.

Looking at Figure S8, in addition to the fact that the *LC*_*50*_ estimates are not very different, their precision also is similar both between option 1 and option 2, and between the use of GUTS-RED-SD and GUTS-RED-IT. This result is quite counterintuitive with the fact that option 1 provides more precise parameter estimates. An explanation may be that the *h*_*b*_ parameter is poorly correlated to the other three parameters. Consequently, in terms of ERA, if only *LC*_*50*_ estimates are needed to decide, estimating *h*_*b*_ or not does not matter a lot.

## Conclusion

Our study clearly illustrated that background mortality (*h*_*b*_) values were almost always different between options 1 and 2. When estimated (option 1), parameter *h*_*b*_ was always closer to the expected background mortality, and its estimate was quite precise. On the other hand, when *h*_*b*_ was calculated from control data (option 2), it was always underestimated. We also noticed that option 2 tended to provide underestimated no-effect concentrations: parameter *z*_*w*_ in GUTS-RED-SD or parameter *m*_*w*_ in GUTS-RED-IT. Parameter *b*_*w*_ displayed a similar pattern for GUTS-RED-SD, meaning that option 2 underestimated the intensity of the chemical effect on individuals. Regarding *k*_*d*_ estimates (i.e., the kinetic quickness), there was no difference between options 1 and 2. Nevertheless, *k*_*d*_ estimates differed according to the GUTS-RED variant, sometimes underestimated, either with GUTS-RED-SD or GUTS-RED-IT. This confirms our inability to anticipate which of the GUTS-RED variants would be more appropriate for a particular species-compound combination. Furthermore, we showed that neither the GUTS-RED variant nor the choice between option 1 and 2 had an impact on any of the goodness-of-fit criteria. Estimating *h*_*b*_ or not only had tiny effects either on *LC*_*50*_ medians or its precision, while the choice of the GUTS-RED variant could sometimes provide different *LC*_*50*_ estimates. Altogether, for ERA mainly needing precise (and accurate) *LC*_*50*_ estimates, estimating the background mortality or not is of little importance. In contrast, benefiting of the most precise parameter estimates for GUTS-RED models may be of strong interest if the objective is to safely predict a survival probability under an environmentally realistic scenarios along which the exposure concentration is varying over time.

## CRediT author statement

**Julie Plantade:** Conceptualization, Data curation, Investigation, Methodology, Validity tests, Visualization, Writing - Original draft preparation. **Virgile Baudrot**: Methodology, R code writing - Reviewing and Editing. **Sandrine Charles:** Investigation, Supervision, Validation, Writing - Reviewing and Editing.

## Acknowledgments

The authors specially address thanks to Professor Xavier Charpentier allowing Julie Plantade to efficiently contribute to this manuscript while conducting her PhD thesis under his supervision. The authors also thank very much Professor Andreas Focks for fruitful exchanges at the beginning of this work in 2020 while we were all confined and isolated each behind our screen. This research work was performed during Julie Plantade’s master internship and did not receive any specific grant from funding agencies in the public, commercial, or not-for-profit sectors.

## Supplementary material and additional information

- *Supplementary material:* All supplementary material is available at https://doi.org/10.5281/zenodo.7565504.
- *Additional information:* All data analyses performed to get the results of this work can identically be reproduced either with the MOSAIC web platform and its GUTS-fit module [18], or directly within the R software and the use of the ‘morse’ package [19].

